# Tissue-specific transcriptomics reveal functional differences in maize floral development

**DOI:** 10.1101/2021.05.09.443315

**Authors:** Hailong Yang, Kate Nukunya, Queying Ding, Beth E. Thompson

**Author notes:** Corresponding author: Beth E. Thompson, Ph.D. Associate Professor, Department of Biology, East Carolina University, Greenville, NC 27858, Email address of Author for Contact. **Author Contributions:** K.N., Q.D, H.Y. and B.T designed the research; K.N., Q.D., and H.Y. performed experiments and analyzed data with B.T. H.Y. and B.T. wrote the manuscript.

## Abstract

Flowers are produced by floral meristems, groups of stem cells that give rise to floral organs. In grasses, including the major cereal crops, flowers (florets) are contained in spikelets, which contain one to many florets, depending on the species. Importantly, not all grass florets are developmentally equivalent, and one or more florets are often sterile or abort in each spikelet. Members of the Andropogoneae tribe, including maize, produce spikelets with two florets; the upper and lower florets are usually dimorphic and the lower floret greatly reduced compared to the upper floret. In maize ears, early development appears identical in both florets but the lower floret ultimately aborts. To gain insight into the functional differences between florets of different fates, we used laser capture microdissection coupled with RNA-seq to globally examine gene expression in upper and lower floral meristems in maize. Differentially expressed genes were involved in hormone regulation, cell wall, sugar and energy homeostasis. Furthermore, cell wall modifications and sugar accumulation differed between the upper and lower florets. Finally, we identified a novel boundary domain between upper and lower florets, which we hypothesize is important for floral meristem activity. We propose a model in which growth is suppressed in the lower floret by limiting sugar availability and upregulating genes involved in growth repression. This growth repression module may also regulate floret fertility in other grasses and potentially be modulated to engineer more productive cereal crops.

**One sentence summary:** Floret-specific differences in cell wall composition and sugar accumulation likely contribute to growth suppression in the lower floret of maize spikelets.

## Introduction

Flowers are essential for plant reproduction and also form fruits and seeds, which are consumed as food. Flowers are produced by floral meristems, undifferentiated groups of stems cells that generate floral organs (Bartlett and Thompson, 2014). Grass flowers (florets) are contained in spikelets, which contain two bracts (glumes) and one to many florets depending on the species. Like other grass flowers, maize florets are highly derived structures. Within two enclosing organs, the lemma and palea, maize flowers contain two lodicules (homologous to petals), three stamens and three carpels, two of which fuse to form the silk (Figure 1).

**Figure 1.**
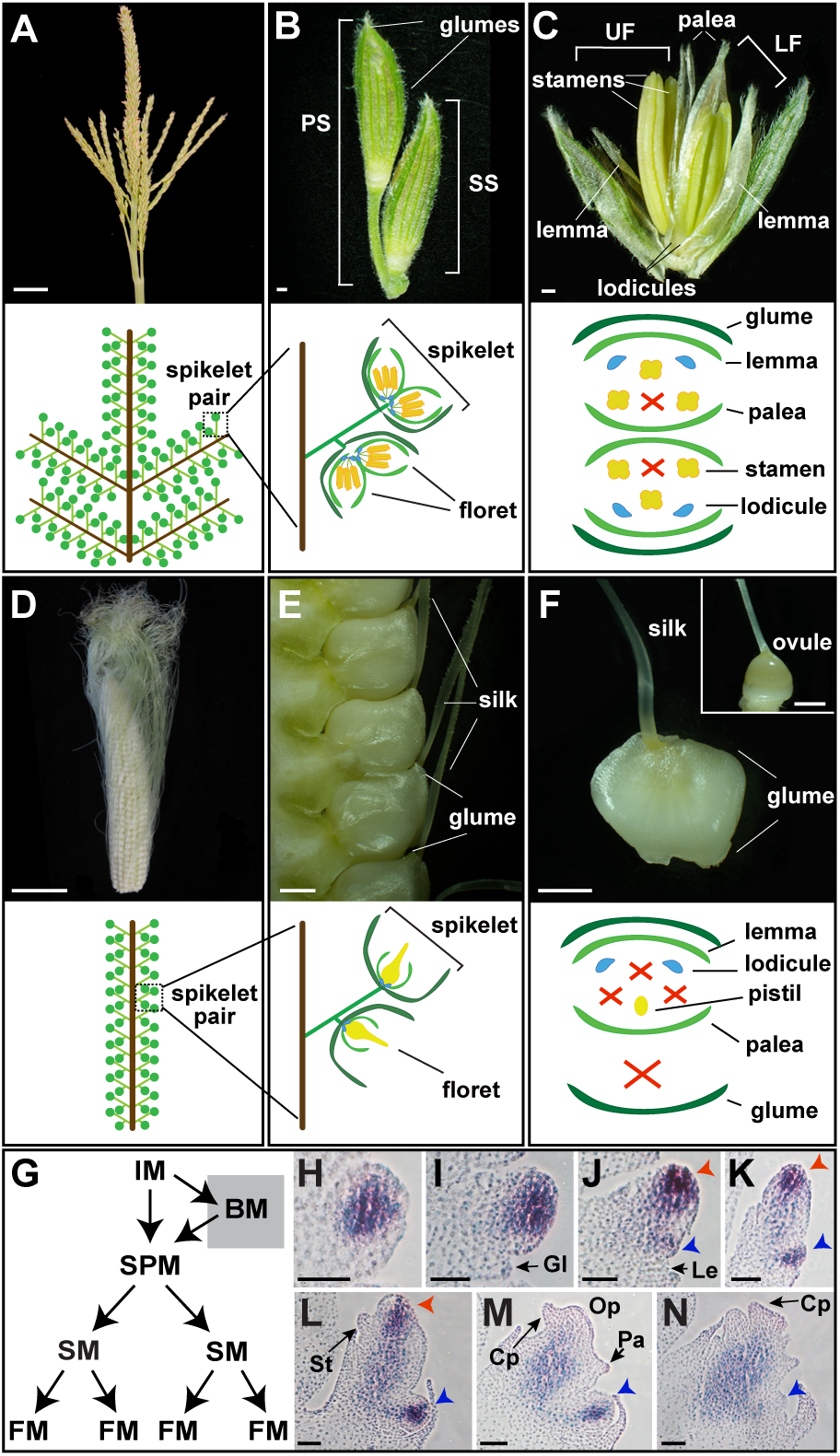
Normal maize floral development. (A) Mature tassel, the male inflorescence. (B) Pair of tassel spikelets. (C) Dissected tassel spikelet, exposing two male florets. (D) Mature ear, the female inflorescence. (E) Mature ear spikelets. (F) Dissected ear spikelet, containing a single female floret. Inset is a mature ovule with glumes and other floral organs removed. (G) Diagram depicting meristems in the inflorescence. (H-N) RNA *in situ* hybridization of the meristem marker, *kn1*, in developing ear spikelets; Red and blue arrowheads indicated upper and lower floral meristems, respectively. PS, pedicellate spikelet; SS, sessile spikelet; IM, inflorescence meristem; BM, branch meristem; SPM, spikelet pair meristem; SM, spikelet meristem; FM, floral meristem; Gl, glume; Op, ovule primordia; Cp, carpel primordia; Lo, lodicule; St, stamen; Le, lemma. Scale bars: (A, D) = 5 cm, (B, C, E, F) = 500 μm, (H-N) = 50 μm.

Maize produces two inflorescences, the tassel and ear, which produce male and female flowers, respectively (Cheng et al., 1983). Unlike Arabidopsis, in which the inflorescence meristem directly initiates floral meristems (FM) on its flanks, grass inflorescence meristems produce a series of higher order meristems before initiating FM. Upon the transition to flowering, the shoot apical meristem (tassel) or an axillary meristem (ear) transitions to an indeterminate inflorescence meristem; inflorescence meristems initiates ordered rows of spikelet pair meristems, which in turn gives rise to two spikelet meristems (Figure 1G) (Thompson and Hake, 2009; Whipple, 2017). The spikelet meristem first initiates the proximal/lower FM (LFM) in the axil of a lemma on the abaxial side of the spikelet meristem (Figure 1, H-L). The origin of the distal/upper FM (UFM) is less clear; one model proposes that the UFM is also initiated as an axillary meristem by the spikelet meristem whereas the second model proposes the spikelet meristem is itself converted to the UFM (Irish, 1997; Chuck et al., 1998).

Both the tassel and ear initiate bisexual flowers and early floral development is very similar in upper and lowers florets (Irish and Nelson, 1989). Carpels abort via programmed cell death in the tassel and stamens arrest shortly after anther formation in the ear (Cheng et al., 1983). In the ear, lower floret abortion is initiated by programmed cell death in the carpel, similar to the carpel abortion program in the tassel (Cheng et al., 1983). Thus, mature ear spikelets contain a single female floret, whereas mature tassel spikelets contain two male florets (Figure 1, A-F).

Spikelets that contain sterile or aborted florets is a common feature of grasses, including cereal crops. In some species, (e.g. maize and barley), floret abortion/sterility is genetically preprogrammed and invariable between individuals whereas in other species (e.g. wheat), the number of aborted florets in a spikelet is variable and influenced by the environment. A few regulators of floral abortion/sterility have been identified (i.e. jasmonic acid in maize, *vrs* genes in barley, and *GN1* in wheat); however, the importance of floral abortion/sterility is still unknown and we know very little about the processes downstream of these high level regulators (Sakuma and Schnurbusch, 2020). To gain insight into the functional differences between florets with different developmental fates, we used laser capture microdissection (LCM) coupled with RNA-seq to globally survey gene expression in UFM and LFM.

## Results

### Upper and lower floral meristems distinct gene expression profiles

Gene expression is dynamic during floral development and to ensure we isolated upper and lower FM at equivalent developmental stages, we isolated FM after initiation of lemma primordia, but before stamen primordia (Figure 2, A-F). LFM development is delayed relative to the UFM (Cheng et al., 1983), and therefore LFM were dissected from older spikelets than UFM. Principal component analysis (PCA) and Pearson’s correlation analysis confirmed UFM and LFM biological samples clustered together and had high reproducibility (Supplemental Figure 1). Approximately 700 genes were differentially expressed between UFM and LFM (238 UFM-enriched, 456 LFM-enriched; FC ≥2 and q <0.05; Figure 2G and Supplemental File 1). Importantly, our data included three UFM-enriched genes with known RNA expression patterns (Supplemental Figure 2 and Supplemental File 2). *zmm8*/GRMZM2G102161 and *zmm14*/GRMZM2G099522 encode MADS-box transcription factors that are broadly expressed in the meristem and floral organs of the upper floret, but not detected in the lower floret (Cacharron et al., 1999; Du et al., 2021). *barren stalk1 (ba1)*/GRMZM2G397518 encodes a bHLH protein required for axillary meristem initiation and is expressed in a diffuse pattern in UFM and in a group of cells at the UFM/LFM boundary, but not detected in LFM (Gallavotti et al., 2004).

**Figure 2.**
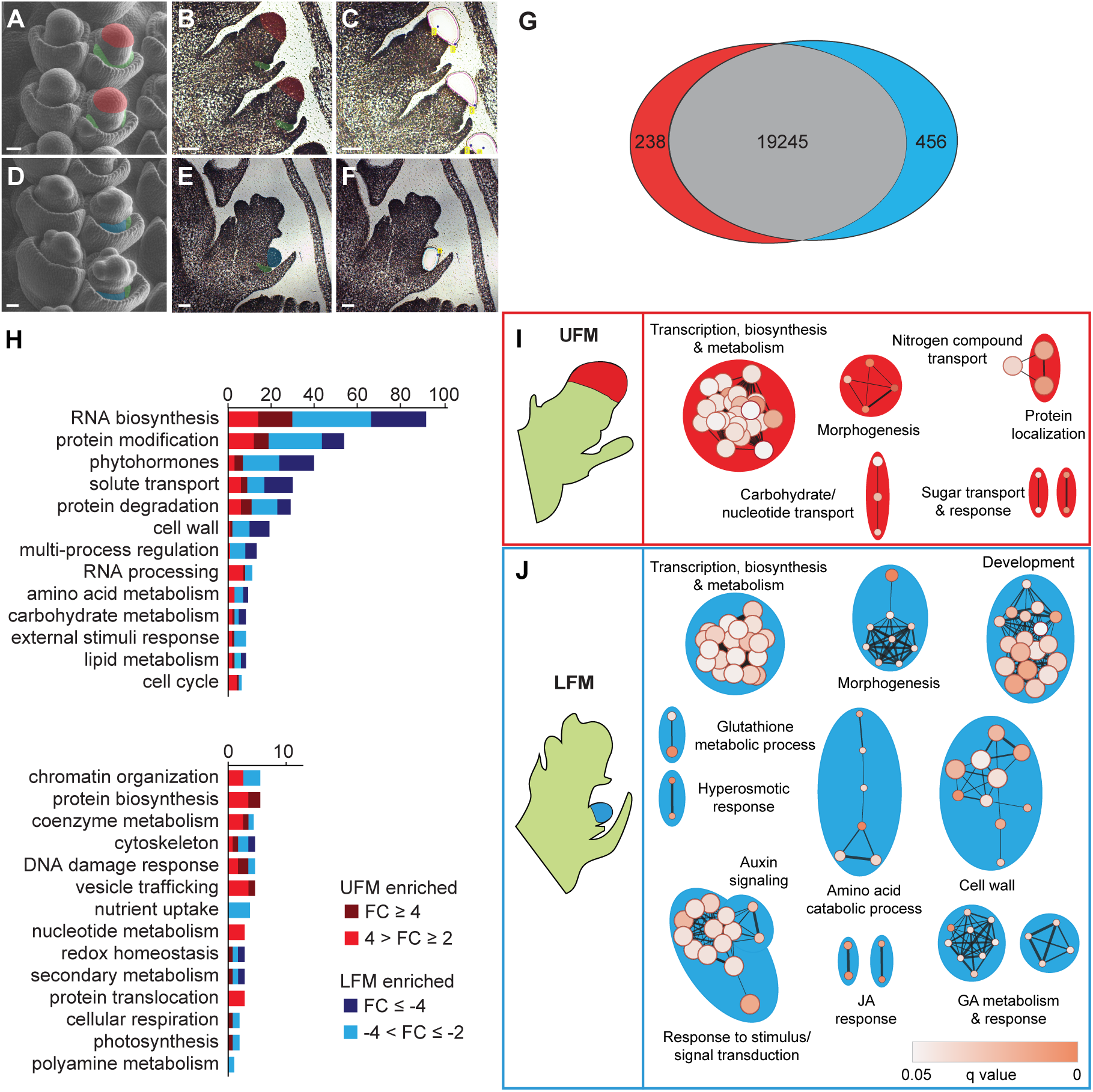
Maize upper and lower floral meristems are enriched for genes belonging to distinct functional groups. SEMs of spikelets at developmental stages of UFM (A) and LFM (D) dissections. Representative images of FM before (B, E) and after (C, F) LCM. False coloring indicates UFM (red), LFM (blue) and lemma primordia (green). (G) Venn diagram depicting DEG (q < 0.05 and fold change ≥ 2) in UFM (red) and LFM (blue). (H) Distribution of DEG in Mapman-annotated functional groups. Genes unassigned to a functional group (99 UFM-enriched and 227 LFM-enriched) are not shown. (I, J) GO enrichment maps for UFM and LFM DEG. Nodes (circles) indicate significantly enriched GO terms. Node size is proportional to number of DEG in each node. Node color indicates statistical significance. Edges (lines) link similar GO terms. Edge thickness is proportional to the number of DEG shared between GO terms. Node clusters were manually labeled based on corresponding GO terms in each cluster. Scale bars = 50 μm.

To gain insight into the biological function of DEG, we predicted gene function using Mapman (Schwacke et al., 2019), gene ontology (GO) enrichment analysis (Gene Ontology Consortium, 2015), and CornCyc, which predicts metabolic pathways (Schläpfer et al., 2017) (Figure 2, H-J and Supplemental File 2). To facilitate the interpretation of hierarchical GO enrichment groups, we used the Cytoscape plug-in, Enrichment Map, to construct functional GO networks (Merico et al., 2010). In general, the UFM was enriched for genes in functional groups associated with growth and primary metabolism, including RNA synthesis and processing, protein synthesis, vesicle trafficking, nucleotide metabolism, and sugar response and transport (Figure 2, H and I; Supplemental File 2). In contrast, the LFM was enriched for genes in functional groups associated with secondary metabolism and dormancy, including phytohormones, protein degradation, amino acid catabolism, and cell wall-related genes (Figure 2, H-J and Supplemental File 2).

We further examined select functional groups to gain insight into the functional patterns of DEG (Supplemental Figure 2-4 and Supplemental File 2). DEG in the RNA biosynthesis group contains several classes of transcription factors (TFs) with well-known roles in plant growth and development, including APETALA2/ETHYLENE RESPONSIVE FACTOR (AP2/ERF), myeloblastosis (MYB), homeobox, basic helix-loop-helix (bHLH), TEOSINTE BRANCHED1/CYCLOIDEA/PROLIFERATING CELL NUCLEAR ANTIGEN FACTOR (TCP), and WRKY TFs (**Supplemental Figure 2**). DEG also included TFs with known functions in maize floral development, including *ba1*/GRMZM2G397518, (Gallavotti et al., 2004), *branched silkless1 (bd1)*/GRMZM2G307119 (Chuck et al., 2002), *gnarley1 (gn1)*/GRMZM2G452178 (Foster et al., 1999a; Foster et al., 1999b), *teosinte branched1 (tb1)*/AC233950.1_FG002 (Hubbard et al., 2002), *Wavy auricle in blade1 (Wab1)*/*branched angle defective1 (bad1)*/GRMZM2G110242 (Hay and Hake, 2004; Bai et al., 2012; Lewis et al., 2014), *GRF-interacting factor1 (gif1)*/GRMZM2G180246 (Zhang et al., 2018), *zfl2*/GRMZM2G180190 (Bomblies et al., 2003), *zmm8*/GRMZM2G102161 and *zmm14*/GRMZM2G099522 (Du et al., 2021). Thus, transcriptional regulation significantly differs between UFM and LFM.

DEG in the phytohormone group function in metabolism and signaling of multiple hormones, including cytokinin, auxin, gibberellin (GA) and jasmonic acid (JA) (Figure 2, H-J; Supplemental Figure 3; Supplemental File 2). Of the four cytokinin-related genes in our DEG set, two cytokinin biosynthesis genes (*czog1*/GRMZM2G168474, GRMZM2G008726) were UFM-enriched and two A-type ARR negative regulators of cytokinin signaling (*crr2*/GRMZM2G392101, GRMZM2G179827) were LFM-enriched, suggesting that cytokinin levels and signaling are higher in UFM relative to LFM. In contrast, auxin, GA and JA-related genes were predominantly enriched in LFM. JA is required for lower floret abortion in the ear (DeLong et al., 1993; Acosta et al., 2009; Lunde et al., 2019; Wang et al., 2020) and three JA biosynthesis genes were LFM-enriched (*lox9*/GRMZM2G017616; *tasselseed1 (ts1)*/GRMZM2G104843; GRMZM2G168404). Seven auxin-related DEGs were enriched in LFM and functioned in auxin synthesis (*tar2*/GRMZM2G066345), transport (*pin3*/GRMZM2G149184; GRMZM2G085236; GRMZM2G037386) and signaling (*aas8*/GRMZM2G053338; *iaa37*/GRMZM2G359924; *bif4*/GRMZM5G864847). GA-related DEG were also LFM-enriched and function in the GA synthesis (*ga20ox1*/AC203966.5_FG005), inactivation (*ga2ox3*/GRMZM2G022679; *ga2ox9*/GRMZM2G152354), and signaling (*gras46*/GRMZM2G001426; GRMZM2G040278; GRMZM2G440543). The LFM was also enriched for three genes encoding Gibberellic Acid Stimulated Arabidopsis (GASA) cysteine-rich polypeptides (*gsl1*/GRMZM2G062527; GRMZM2G077845; GRMZM2G150688), which in Arabidopsis, are induced by GA and have broad functions in defense and development (Roxrud et al., 2007; Zhong et al., 2015). These gene expression profiles suggest that hormone accumulation and signaling differs between UFM and LFM of maize ears, with high cytokinin in the upper floret and high auxin, GA and JA in the lower floret.

We investigated the spatial expression patterns of 10 DEG by RNA *in situ* hybridization (Figure 3). DEG had dynamic and specific expression patterns in developing spikelets, suggesting they have key roles in floral development. AC217050.4_FG006 (encodes a 14-3-3 protein, log_2_FC = 1.124), *AP2/EREBP transcription factor 26 (ereb26)*/GRMZM2G317160 (log_2_FC = -1.162), and *chromatin complex subunit A 101 (chr101)*/GRMZM2G177165 (log_2_FC = -1.251) were broadly expressed in both upper and lower FM (Figure 3, A-C); AC217050.4_FG006 and *chr101*/GRMZM2G177165 were also present in stamen and carpel primordia (Figure 3, A and C; Supplemental Figure 5). As previously shown, *gif1*/GRMZM2G180246 (log_2_FC = 1.063) was expressed in a ring around developing UFM and at the base of palea in upper florets (Zhang et al., 2018), and showed a similar expression pattern in lower florets (Figure 3D). GRMZM2G101682 (grass-specific gene of unknown function, log_2_FC = -1.009) was also expressed in both UFM and LFM (Figure 3E), with strong expression restricted to the outermost cell layer. We also observed expression in spikelet meristem, stamen and carpel primordia (Figure 3E and Supplemental Figure 5). In developing shoots, GRMZM2G101682 is localized to the L1 layer of boundary regions between initiating organs and the preligular band of developing leaves, but is not expressed in the mersitem itself (Johnston et al., 2014). *Histone H1-like*/GRMZM2G069911 (log_2_FC = -1.233) and GRMZM2G180870 (*XYLOGLUCAN ENDOTRANSGLUCOSYLASE/HYDROLASE 9* (*XTH9)* homolog, log_2_FC = -1.059) were expressed in punctate patterns characteristic of genes involved in cell division (Figure 3, F and G; Supplemental Figure 6) (Asai et al., 2002; Kim et al., 2007; Ikeda-Kawakatsu et al., 2009). In the SAM, stem cells at the tip of the meristem have lower cell division rates compared to cells in axillary primordia (Satterlee et al., 2020). If stem cells at the tips of FM also have lower cell division rates, the apparent LFM-enrichment of cell division genes likely reflects the higher proportion of “tip stem cells” (with lower cell division rates) in UFM samples relative to LFM samples. Finally, three genes were localized to a unique boundary region between the upper and lower florets. GRMZM2G114552 (log_2_FC = 2.616) encodes a Bowman-Birk type trypsin inhibitor (*BBTI*) and was expressed in a discrete domain on the abaxial side of UFM but not detectable in LFM (Figure 3H). *BBTI* was also expressed at the boundaries of initiating spikelet pair and spikelet meristems, and at the base of palea in the upper floret (Supplemental Figure 7). A *pectate lyase* homolog, GRMZM2G131912 (log_2_FC = -1.281), and *arginine decarboxylase1 (adc1)*/GRMZM2G396553 (log_2_FC = -1.216) were present in discrete domains on the adaxial side of the LFM at the boundary with the upper floret but not in the upper floret itself (Figure 3, I and J; Supplemental Figure 7). *BBTI, adc1,* and *pectate lyase* were also expressed in this boundary region in tassel florets (Supplemental Figure 7), indicating that this boundary expression is not unique to the ear.

**Figure 3.**
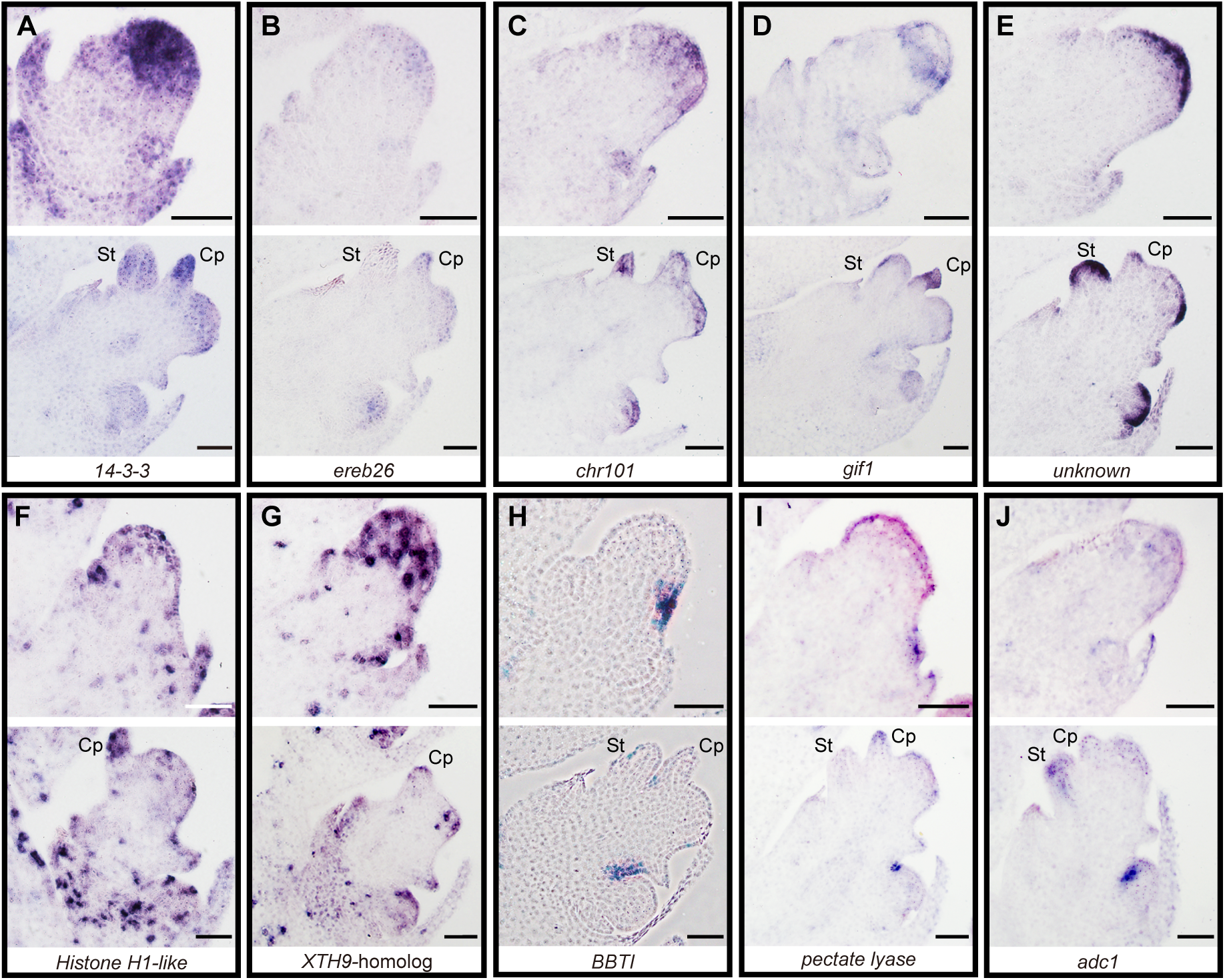
UFM and LFM DEG have distinct RNA expression patterns. (A) AC217050.4_FG006, 14-3-3 protein. (B) GRMZM2G317160/*ereb26*. (C) GRMZM2G177165/*chr101*. (D) GRMZM2G180246/*gif1*. (E) GRMZM2G101682, unknown function. (F) GRMZM2G069911, *histone H1-like*. (G) GRMZM2G180870, *AtXTH9 (XYLOGLUCAN ENDOTRANSGLUCOSYLASE/HYDROLASE 9)* homolog. (H) GRMZM2G114552/*BBTI*, Bowman-Birk type (proteinase/bran trypsin) inhibitor. (I) GRMZM2G131912, *pectate lyase* homolog. (J) GRMZM2G396553/*adc1*. Developmental stages at which UFM (top) and LFM (bottom) were dissected are shown for each gene. St: Stamen; Cp: carpel primordium. Scale bars = 50 μm.

### Pectin modification is dynamic during spikelet development and differs between upper and lower florets

GO and Mapman functional analyses indicated that DEG belonged to multiple functional groups (Figure 2), several of which made sense based on their well-established roles in development (i.e. transcription, development, morphogenesis, hormones); however, other functional groups were more surprising. We were particularly intrigued by enrichment of cell wall-related genes in LFM and sugar-related genes in UFM, and thus further investigated these functional groups.

Our DEG set included 20 Mapman-annotated cell wall-related genes, 18 of which were enriched in LFM (Figure 4). Indeed, RNA *in situ* hybridization confirmed that GRMZM2G131912 (*pectate lyase* homolog), is exclusively expressed in the lower floret, adjacent to the UFM/LFM boundary (Figure 3I). Cell wall-related DEG were involved in synthesis or modification of all major cell wall components, including cellulose (one gene), lignin (two genes), hemicellulose (four genes), and pectin (five genes), as well as arabinogalactan proteins (six genes) and expansins (two genes) (Figure 4A and Supplemental File 2). Most of these genes are involved in synthesis and modification of the primary cell wall, which is synthesized and continuously deposited around dividing or expanding cells, including meristems (Cavalier et al., 2008; Keegstra, 2010; Sampathkumar et al., 2019).

**Figure 4.**
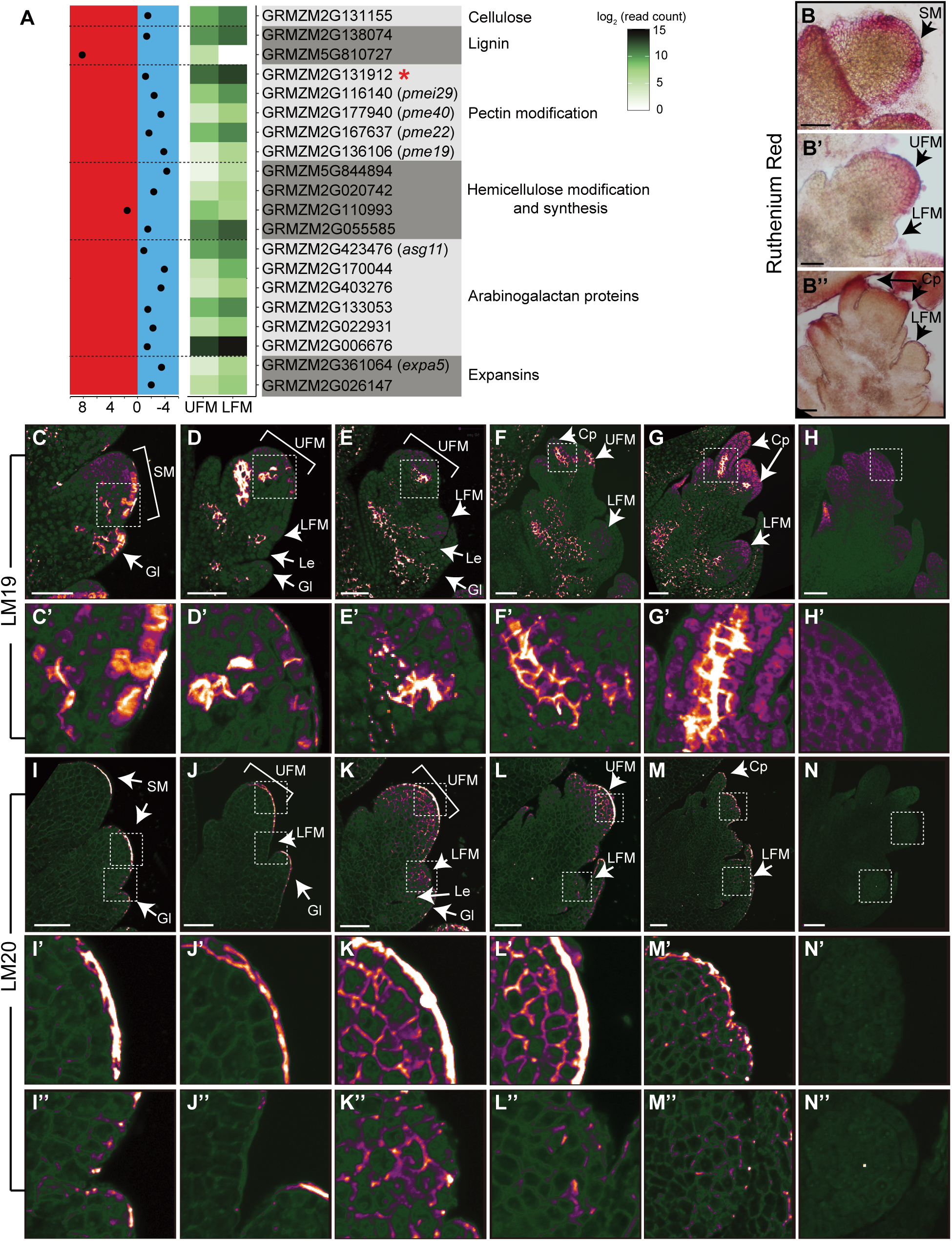
The UFM and LFM have distinct cell wall compositions. (A) Expression profiles for DEG in the MapMan Cell Wall functional group. Left panel indicates log_2_ (fold change) for each gene. UFM-enriched genes are plotted on the left (red) and LFM-enriched genes on the right (blue). Middle panel shows an expression heatmap. Red * indicates gene with known RNA *in situ* hybridization patterns. Ruthenium red staining of acidic pectin in SM (B), early (B’) and late (B’’) ear florets. (C-G) LM19 immunostaining of low methylesterified HG. (H) Negative control lacking primary antibody to show background autofluorescence using the same laser settings as (C-G). White boxes indicate zoomed in areas shown in (C’-H’). (I-M) LM20 immunostaining of high methylesterified HG in developing spikelets. (N) Negative control lacking primary antibody to show background autofluorescence using the same laser settings as (I-M). White boxes indicated zoomed in areas corresponding to UFM (I’-N’) and LFM (I’’-N’’). Weak, diffuse cytoplasmic signal in (C-N) is background autofluorescence, which varies due to incomplete quenching. Micrographs were false colored using ImageJ (Orange Blue icb look-up table) to visualize signal intensity. Scale bars = 50 μm.

Differential expression of cell wall-related genes suggested that UFM and LFM have different cell wall compositions and/or modifications. Therefore, we stained the major cell wall components in developing spikelets, including cellulose (calcofluor white/fluorescent brightener 28), lignin (phloroglucinol-HCl), and pectin (ruthenium red). To confirm our staining protocols accurately reflected cell wall composition, we first stained vasculature tissue, where cell wall composition is well-characterized (Supplemental Figure 8) (Chen et al., 2006; Verhertbruggen et al., 2009; Pesquet et al., 2013; Hu et al., 2017; Torode et al., 2018). Lignin is predominantly found in secondary cell walls, which only form after cells have stopped expansion (Zhong et al., 2019). As expected for meristematic tissue, lignin staining was weak or undetectable in inflorescence primordia (Supplemental Figure 8), with no indication of floret-specific accumulation. Cellulose accumulated at the periphery of all cells and appeared similar in both florets (Supplemental Figure 8). We visualized pectin using ruthenium red, which preferentially stains acidic pectin (Ruzin, 1999), and observed striking differences in pectin distribution between UFM and LFM. Ruthenium red strongly stained the L1 layer of spikelet meristem and glume primordia; staining persisted in the L1 of UFM, however was much weaker or absent in LFM (Figure 4B).

To examine pectin composition in more detail, we used two two monoclonal antibodies, LM19 and LM20, which recognize the low methylesterified (acidic) and high methylesterified forms of homogalacturonan (HG), the most abundant pectic polysaccharide (Verhertbruggen et al., 2009). We observed dynamic LM19 staining during floral development, suggesting that HG methylesterification is developmentally regulated (Figure 4, C-H). LM19 (low methylesterification) strongly stained the L1 layer of spikelet meristem and glume primordia, similar to ruthenium red staining (Figure 4C). LM19 also stained cells at the base of glumes, and strongly stained incipient and initiating floral organ primordia (Figure 4, D-G). We observed weak staining at the base of the LFM, and occasionally observed a couple of brightly staining cells that appeared to correspond to palea primordia (Figure 4G). However, we never observed strong LM19 staining associated with organ primordia in LFM.

LM20 (high methylesterified HG) showed a dramatically different staining pattern than LM19 (Figure 4, I-N). LM20 weakly stained the periphery of most or all cells in developing spikelets and intensely stained the apical surface of the L1 layer of spikelet meristem and young UFM (Figure 4, I-K). LM20 staining persisted throughout UFM development, but its localization became more punctate and primarily accumulated at cell junctions (Figure 4, K-M). In LFM, LM20 also stained cell peripheries, often forming puncta at cell junctions (Figure 4, J-M). Although variable, we never observed strong LM20 staining in LFM as we did in UFM. These results indicate that pectin is dynamically regulated during floral development and pectin modification differs between upper and lower FM. Furthermore, demethylesterified pectin strongly accumulates in incipient and initiating organ primordia of UFM, indicating pectin may also play a role in organ initiation in maize.

### Sugar-related genes and starch are differentially regulated in upper and lower floral meristems

The ability to coordinate energy and carbon availability with plant growth is critical. Hexose sugars generated through photosynthesis in source tissues are converted to sucrose for transport to sink tissues and starch for storage. Carbohydrates are required not only to provide chemical energy required for plant growth, but also to generate nucleotides and construct cell walls around newly divided and expanding cells (Sampathkumar et al., 2019). GO and MapMan analysis indicated UFM were enriched for genes involved in carbohydrate/sugar transport and response (Figure 2, H and I; Supplemental File 2), while the LFM was enriched for multiple members of the SNF1-related protein kinase 1 (SnRK1) signaling pathway (Figure 5, A and B).

**Figure 5.**
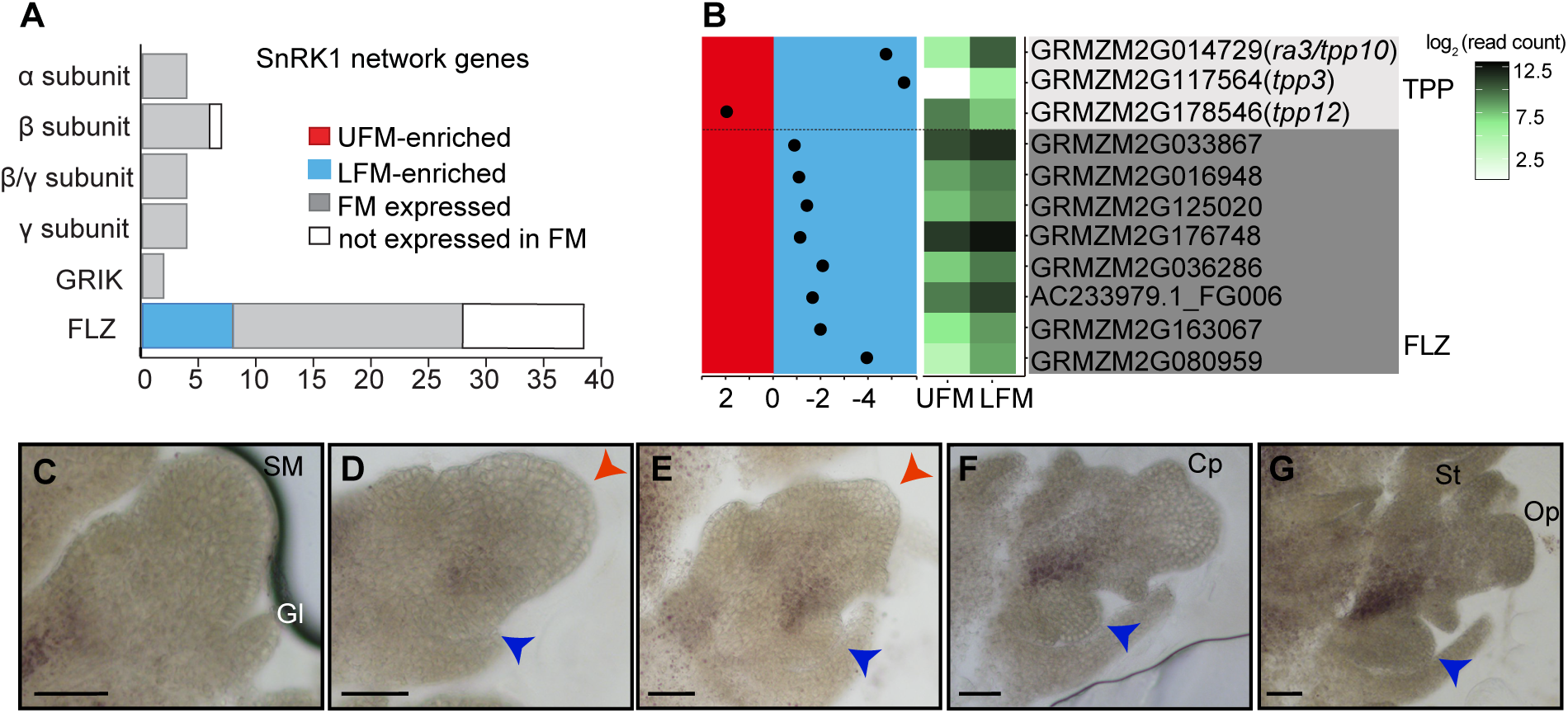
Sugar metabolism likely differs in UFM and LFM. (A) Summary of predicted SnRK1 signaling network genes expressed in FM samples. (B) Expression profiles for sugar-related DEG; layout is the same as cell wall-related genes in Fig 4. (C-G) Starch accumulates at the boundary between the UFM and LFM in developing spikelets. Red and blue arrowheads indicate UFM and LFM respectively. Scale bars = 50 μm.

The SnRK1 pathway is a key regulator of plant growth and energy homeostasis (Baena-González et al., 2007) and is regulated both by trehalose-6-phosphate (T6P) levels (Baena-González and Lunn, 2020) and interactions with FCS-like zinc finger (FLZ) proteins (Nietzsche et al., 2014; Jamsheer K et al., 2018b; K et al., 2019). Trehalose is present in trace amounts in plants and its primary role is likely in sugar sensing and signaling, rather than chemical energy storage (Wingler, 2002; Figueroa and Lunn, 2016). The trehalose precursor, T6P, is synthesized from UDP-glucose and glucose-6-P by T6P synthase (TPS) and thought to be the active signaling molecule; T6P is converted to trehalose by trehalose-6-phosphate phosphatases (TPPs) (Cabib and Leloir, 1958). T6P levels are positively correlated with sucrose levels and T6P sensing may be a key mechanism by which plants monitor sucrose availability (Figueroa and Lunn, 2016). The maize genome contains 13 TPP genes, of which *ramosa3 (ra3)/tpp10/*GRMZM2G014729 is the most extensively studied and is required for spikelet pair and spikelet meristem determinacy in the inflorescence (Satoh-Nagasawa et al., 2006).

Interestingly, *ra3* promotes meristem determinacy at least in part independent of its enzymatic activity and likely functions in transcriptional regulation (Claeys et al., 2019a; Demesa-Arevalo et al., 2021). Both *ra3* and *tpp3*/GRMZM2G117564 were LFM enriched, suggesting that sugar signaling could be critical in the LFM. Recently work also showed that *ra3* functions redundantly with the HD-ZIP TF, *grassy tillers1 (gt1)* to repress carpel growth in tassel florets. Particularly relevant to this work, the lower floret fails to abort in *ra3; gt1* double mutants, demonstrating that our approach identified genes with functional differences in the upper and lower florets (Klein et al, in preparation). The plant-specific *FLZ* gene family is defined by the presence of a ∼50 amino acid FLZ domain, which interacts with SnRK1 (K and Laxmi, 2014; Jamsheer K et al., 2018b). While the function of many *FLZ* genes is unknown, they have been implicated in ABA, sugar, and energy response in Arabidopsis (K and Laxmi, 2015; Jamsheer K et al., 2018a) and are thought to act as adapters between SnRK1 and other proteins (Tsai and Gazzarrini, 2014; Jamsheer K et al., 2018b; K et al., 2019). Based on MapMan annotations, the maize genome contains 39 *FLZ* genes in the “multiprocess regulation” functional group, 28 of which were expressed in our FM samples. Strikingly, nearly one-third (8/28) of the FM-expressed *FLZ* genes were DE between UFM and LFM, all of which were LFM-enriched (Figure 5 and Supplemental File 2). All core components of the SnRK1 signaling pathway were expressed in our FM samples, but only *FLZ* genes were DE between UFM and LFM (Figure 5A). Thus, the SnRK1 signaling pathway may function in the LFM to sense and respond to low sugar availability.

Determining sugar accumulation and distribution *in situ* is challenging due to a lack of dyes or other mechanisms to detect specific sugars. Most sugar analysis requires grinding tissue and measuring overall sugar levels, which precludes the cellular-level resolution required to detect spatial differences in sugar accumulation in a developing tissue. Starch, the major storage carbohydrate in plants, however, can be easily visualized by iodine staining (Zhang et al., 2019). In the inflorescence, starch accumulates at the base of developing spikelet meristems, but not in the spikelet meristem itself (Figure 5C). After LFM initiation, starch begins to accumulate at the base of UFM, near the boundary with LFM (Figure 5, D and E). Starch accumulation intensifies and becomes more defined at the boundary between UFM and LFM in older spikelets (Figure 5, E-G). Strikingly, we did not observe detectable starch in LFM at any stage of spikelet development (Figure 5, C-G). We observed similar starch accumulation in tassels (Supplemental Figure 8, N-R), indicating starch distribution is carefully regulated in developing spikelets of both male and female inflorescences and sugars accumulate differently in UFM and LFM.

## Discussion

### Upper and lower floral meristems are not developmentally equivalent

Floral meristems are typically regarded as functionally equivalent, regardless of where they are initiated on the plant. In Arabidopsis, for example, all FM form as axillary meristems on the flanks on an inflorescence meristem, produce identical flowers, and appear to have the same developmental potential (Liu et al., 2009). In maize, spikelet meristems are often depicted as initiating two equivalent FM (Figure 1G), however this depiction is misleading and upper and lower FM are likely functionally divergent from the time of initiation. The Andropogoneae tribe, which includes maize along with the key crops sorghum and sugarcane, produce paired spikelets, with two florets per spikelet. Upper and lower florets in the Andropogoneae are typically dimorphic; the upper floret is often hermaphroditic whereas the lower floret is usually reduced or sterile (Le Roux and Kellogg, 1999). Although tassel spikelets produce two morphologically identical staminate florets, ear spikelets, in which the lower floret aborts, are more representative of the Andropogoneae, and findings in the ear may be relevant more broadly to other species in the tribe. Floral abortion and sterility are common in the cereals and the mechanisms that regulate lower floret growth in maize may also apply to other cereal crops.

We sought to understand the functional differences of the upper and lower florets by globally surveying gene expression in the UFM and LFM of maize ears. Both UFM and LFM expressed a broad set of genes, including genes previously implicated in floral development and/or meristem function. Approximately 3.5% of genes are differentially expressed between UFM and LFM (Figure 2G and Supplemental File 1), which is consistent with previous molecular and genetic analyses. At least two maize mutants differentially affect the upper and lower florets. In *bearded-ear (bde)* mutants, UFM are indeterminate whereas LFM initiate additional floral meristems and lose FM fate (Thompson et al., 2009). In *rf2* mutants, stamens arrest in lower, but not upper florets (Liu et al., 2001). Microarray analysis indicates that approximately 9% of genes are differentially expressed in equivalently staged anthers from the upper and lower florets (Skibbe et al., 2008); thus, floret-specific gene expression persists even in differentiated floral organs. Finally, two MADS-box genes, *zmm8* and *zmm14*, are only detectable in the upper florets by RNA *in situ* hybridization (Cacharron et al., 1999; Du et al., 2021). These data support the model that UFM and LFM use distinct gene regulatory networks and have divergent developmental fates from the very earliest stages of floral development.

The divergent developmental fates of UFM and LFM may be due in part to their distinct ontogenies. The LFM is clearly an axillary meristem and associated with formation of auxin maxima and with novel expression of shoot and floral meristem markers, such as *knotted1 (kn1)* and *bde,* respectively (Figure 1, H-N) (Jackson et al., 1994; Gallavotti et al., 2008; Thompson et al., 2009). In contrast, formation of the UFM is not associated with an auxin maximum, which supports the model that the UFM is not an axillary meristem but rather that the spikelet meristem is converted to the UFM (Gallavotti et al., 2008). We found the LFM was enriched for auxin-related genes (Figure 2, H and J; Supplemental Figure 3; Supplemental File 2), which could reflect the axillary meristem identity of LFM, but not UFM.

Transcriptional regulatory networks also differ between upper and lower florets. One of the largest groups of DE genes identified in our analysis were transcriptional regulatory proteins, which included TF classes with key functions in plant development (i.e. TCP, WRKY, homeobox, AP2/ERF). These experiments were motivated in part by the *bde* mutant phenotype, which as previously mentioned, promotes FM determinacy in the upper floret and FM fate in the lower floret. *bde* encodes a MADS-box TF and we hypothesized that these floret-specific phenotypes were caused by disruption of distinct BDE-containing complexes in the UFM and LFM, resulting in misregulation of different target genes (Thompson et al., 2009). Surprisingly, *zmm8* and *zmm14* were the only two DE MADS-box genes identified in our samples (Supplemental Figure 2 and Supplemental File 2), both of which were previously shown to be strongly UFM-enriched and have been hypothesized to act as upper floret selector genes (Cacharron et al., 1999). Recent analysis of *zmm8; zmm14* double mutants, however, indicate that *zmm8/zmm14* promote FM meristem determinacy in both florets and do not have floret-specific functions (Du et al., 2021). Although *zmm8/zmm14* are highly UFM-enriched in our data, they are also expressed in LFM (Supplemental File 1). Combined with the fact that we did not identify clear candidates for lower floret selector genes, these data suggest that upper versus lower floret selector genes may not exist. We favor the hypothesis that the developmental history and anatomy of upper versus lower florets lead to physiological differences between the florets (i.e. energy availability or hormone status), which can cause floret-specific mutant phenotypes and ultimately determine UFM versus LFM fate.

### Genes associated with growth repression are enriched in the lower floral meristem

Plants must be able to alter growth and development in response to both internal and external cues, including energy status. Sugar, mainly in the form of glucose and fructose, is produced by photosynthesis in source tissues and transported as sucrose to sink tissues, such as developing seeds. Once localized to sink tissues, sucrose can be converted to glucose and used for chemical energy, as a structural component of cells (e.g. cell walls) or stored for later use. Sugar is also an important signaling molecule and functions in diverse processes. The lower floret is enriched for genes involved in growth repression, and our data suggests low sugar availability in the lower floret may contribute to this growth repression.

The conserved SnRK1 protein kinase (homologous to yeast Snf1 and animal AMPK1) is a key mechanism by which plants sense nutrient availability and maintain energy homeostasis. SnRK1 stimulates pathways that inhibit growth and increase catabolism in response to energy starvation (Baena-González et al., 2007). SnRK1 senses energy status primarily through repression by T6P, which is a proxy for carbon availability (Smeekens, 2015; Figueroa and Lunn, 2016). Our data suggest a model in which low sugar availability in the lower floret suppresses growth via the SnRK1 signaling pathway (Figure 6). The LFM is enriched for RNAs encoding two TPP enzymes (*ra3/tpp10* and *tpp3*) (Figure 5B), consistent with previous reports that showed *ra3* RNA is localized to the UMF/LFM boundary (Satoh-Nagasawa et al., 2006). Increased TPP levels in the LFM are predicted to decrease T6P levels and thus activate SnRK1; the UFM is enriched for a single *tpp* gene (*tpp12*) (Figure 5B), suggesting that T6P levels may be critical throughout the spikelet. In other developmental contexts, the primary role of RA3 (and other TPPs) appears to be in transcriptional regulation, and not direct modulation of T6P levels (Claeys et al., 2019b; Demesa-Arevalo et al., 2021). Regardless, RA3/TPPs are likely to either directly regulate or respond to sugar levels, suggesting that sugar levels are carefully regulated in the spikelet. Both the UFM and LFM express RNAs corresponding to all core components of the SnRK1 signaling pathway, most of which are not differentially expressed (Figure 5A). The LFM, however, showed enrichment of eight *FLZ* genes (Figure 5, A and B), which likely act as adaptors for SnRK1. Although relatively little is known about *FLZ* genes, *FLZ* RNA levels respond to sugar, hormones and abiotic stress and generally are associated with energy deprivation (K and Laxmi, 2015; Jamsheer K et al., 2018a). In addition to SnRK1 subunits, FLZ interacts with TFs and other developmental regulators, including homologs of LFM-enriched genes (TCP, homeobox TFs, GAI, DELLA) (Nietzsche et al., 2016; K et al., 2019). Thus, LFM-enriched *FLZ* genes likely direct SnRK1 to specific targets that function in floral development. The LFM also showed enrichment of genes involved in protein degradation and amino acid catabolism (Figure 2, H and I; Supplemental Figure 4), consistent with active SnRK1 in LFM. High T6P levels have been correlated with increased expression of genes involved in primary metabolism (Oszvald et al., 2018), and indeed, the UFM was enriched for genes involved in RNA processing and protein biosynthesis (Figure 2H and Supplemental File 2).

**Figure 6.**
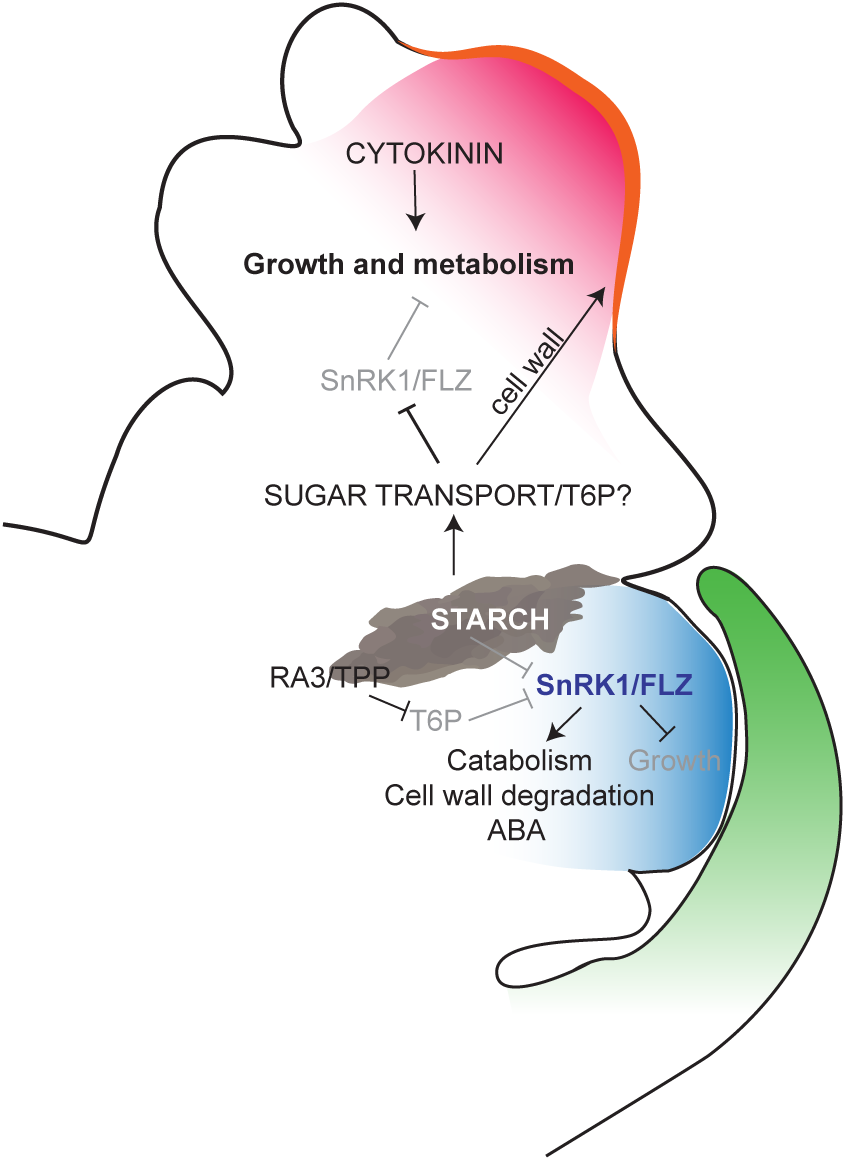
A model for growth regulation in developing florets. In UFM (red), high cytokinin and sugar availability promote growth and metabolism, including synthesis of new cell walls (orange). In LFM (blue), low sugar availability activates the SnRK1/FLZ pathway to repress growth and increase catabolism. Starch accumulates at the UFM/LFM boundary and may supply sugar to the UFM and/or be important for boundary formation.

T6P also promotes starch accumulation, the major storage carbohydrate in plants (Figueroa and Lunn, 2016). Starch gradually accumulates during spikelet development (Figure 5, C-G), presumably increasing the strength of the inflorescence as a sink tissue. Starch does not accumulate throughout the spikelet, but rather accumulates in a defined region at the boundary between the upper and lower floret and appears to be excluded from the lower floret. In animals, the SnRK1 homolog, AMPK1, binds and is negatively regulated by glycogen (Janzen et al., 2018), which is analogous to starch in plants, raising the intriguing possibility that SnRK1 directly binds and is regulated by starch. One possibility is that excluding starch from LFM signals energy starvation and growth repression in the lower floret. Starch accumulation is similar, however, in tassel spikelets (Supplemental Figure 8, N-R) where the lower floret does not abort suggesting that starch distribution may have another function in floral development or that the tassel has another mechanism to overcome starch deprivation.

In addition to its function in sugar signaling and homeostasis within a tissue, T6P can affect sugar utilization and distribution at the whole plant level and is a key regulator of the source/sink balance (Figueroa and Lunn, 2016). For example, low T6P levels in the inflorescence increase sucrose transport to inflorescences and prolonged photosynthesis in leaves (Nuccio et al., 2015; Oszvald et al., 2018). The interaction between source and sink tissues is well-known to affect timing of senescence and many DEG in our data set have been directly or indirectly implicated in senescence. We propose the LFM executes a senescence-like program to repress growth in the lower floret (Figure 6). This model is consistent with previous work showing that the Arabidopsis senescence-inducible promoter, p*SAG12,* is sufficient to drive expression of the cytokinin biosynthesis gene, *isopentenyl transferase1 (ipt1),* in the lower floret and inhibit floret abortion (Young et al., 2004). In our data, the cytokinin biosynthesis gene, GRMZM2G091837 (homologous to rice *lonely guy)* is UFM-enriched and two A-type ARR genes (Supplemental Figure 3 and Supplemental File 2), which are negative regulators of cytokinin signaling are LFM-enriched, consistent with high cytokinin in the upper floret relative to the lower floret. Increased sucrose in wheat also decreases floral abortion, suggesting that sugar signaling and homeostasis is a key regulator of floral abortion in cereal crops (Ghiglione et al. 2008).

Other LFM-enriched genes have been directly or indirectly implicated in senescence. The senescence-inducible chloroplast stay-green protein, GRMZM2G091837 is associated with delayed senescence and UFM-enriched. *Malate synthase 1* (*Mas1/*GRMZM2G102183) and arabidopsis homologs of several LFM-enriched *FLZ* genes are induced in senescing tissues in maize and arabidopsis respectively (Yandeau-Nelson et al., 2011; K and Laxmi, 2015; Jamsheer K et al., 2018a). A recent study by Sekhon et al. (2019) used a combination of GWAS and gene expression analysis to identify 62 genes associated with natural variation of senescence in maize. We noted glutathione S-transferases (GSTs) are both LFM-enriched (5 LFM-enriched and 2-UFM enriched GSTs) and senescence-associated (3 GSTs) (Supplemental Figure S4), and indeed *gst41*/GRZM2G097989 overlapped both lists. In addition to known regulators of senescence (i.e. cytokinin and T6P/SnRK1), the cell wall has emerged as a contributor to senescence (Sekhon et al., 2012; Sekhon et al., 2019). Cell walls can act as secondary sinks for sugar and thus may affect the source/sink balance (Sekhon et al., 2019). Cell wall breakdown may also release hexoses that could be used for energy. While not directly implicated in senescence, we also noted the LFM was enriched for two Rapid Alkalinization Factor (RALF)/RALF-like (RALFL) peptides, which are associated with repressed growth (Blackburn et al., 2020). Thus, the LFM is broadly enriched for genes associated with senescence and growth repression.

Plant growth requires the synthesis of new cell walls as cells divide and modification of existing cell walls to allow for cell expansion. In eudicots, pectin composition and modification is developmentally regulated and has been implicated in multiple aspects of plant growth and development (Saffer, 2018). Pectins are complex galacturonic acid-rich polysaccharides, of which homogalacturonan (HG) is the most abundant (Harholt et al., 2010). HG is deposited in the cell wall in a highly methylesterified form and can be demethylesterified by pectin methylesterseses (PMEs); in Arabidopsis, demethylesterification of HG regulates primordia initiation and phyllotaxy (Peaucelle et al., 2008; Peaucelle et al., 2011). Grass cell walls contain significantly less pectin than eudicot cells walls (∼5% in grasses vs 20-35 % in eudicots) (Vogel, 2008) and what role, if any, pectin modification plays in primordia initiation and phyllotaxy in grasses is unclear. Our data show that demethylesterified pectin is clearly associated with floral organ primordia initiation in the UFM (Figure 4, C-H), strongly suggesting that pectin’s role in organ initiation is conserved in grasses. We did not see similar accumulation associated with organ primordia in the lower floret, which could be reflective of general growth repression in the lower floret.

### Does the boundary between upper and lower florets affect floral meristem activity?

Boundary regions between meristems and initiating primordia are essential to separate groups of cells with different developmental fates and can also affect the activity of adjacent cell populations (Wang et al., 2016; Richardson and Hake, 2018). In grass inflorescences, meristem determinacy is controlled by groups of genes expressed in regions adjacent to the meristem that form boundaries with organ primordia. For example, the *ramosa* regulatory module is expressed at the base of and limits spikelet pair meristem determinacy (Eveland et al., 2014); similarly, *bd1* and *indeterminate spikelet1* (*ids1*) TFs are expressed at the base of and limit spikelet determinacy (Chuck et al., 1998; Chuck et al., 2002). In barley *compositum1* (*com1*) mutants, determinate spikelets on the main rachis are transformed into indeterminate branches due to defective boundary formation. The *ra3* ortholog, along with other sugar and cell wall-related genes are misexpressed in *com1* mutants, suggesting that sugar-signaling and cell wall changes are critical for boundary formation (Poursarebani et al., 2020). Thus, boundary regions adjacent to meristems may function as novel signaling centers that regulate meristem activity (Whipple, 2017), although the mechanism is unclear.

Our data suggest that a similar boundary program may function at the UFM/LFM boundary. First, *ra3* RNA is localized to the boundary (Satoh-Nagasawa et al., 2006) and overlaps with starch accumulation (Figure 5, C-G). We also identified the *com1* ortholog, *Wab1/bad1,* as LFM-enriched gene in our samples (Supplemental Figure 2 and Supplemental File 2). Because of the small size of LFM, we likely isolated boundary genes in our LFM samples that were excluded from UFM samples in which we were able to isolate the “tips” of the meristems. Our data also suggest cell walls are differentially regulated in UFM and LFM (Figure 4). Indeed, *pectate lyase* was localized to a discrete domain at the UFM/LFM boundary (Figure 3I and Supplemental Figure 7) and boundary regions are often characterized by stiffer cell walls (Richardson and Hake, 2018). *Arginine decarboxylase1,* a key enzyme required for synthesis of the polyamines, is also expressed at the UFM/LFM boundary (Figure 3J and Supplemental Figure 7). Polyamines have diverse functions in plants ranging from stress responses to growth and development, including flower bud formation (Chen et al., 2018). *ba1* is also expressed in the domain at the UFM/LFM boundary and later below the palea (Gallavotti et al., 2004). Finally, *BBTI* is expressed in the upper floret at the UFM/LFM boundary and at the base of the palea (Figure 3H and Supplemental Figure 7). The palea expression of *BBTI* and *ba1* is particularly intriguing; in barely, *com1* is also expressed in palea and *com1* mutants have enlarged palea cells with thinner cell walls (Poursarebani et al., 2020). Thus, palea may also have important boundary functions or alternatively, boundary regulatory modules may be redeployed in palea development.

Understanding the genetic and physiological processes that regulate floret abortion and sterility is a necessary first step to engineer maize and other cereal crops with increased floret fertility. Our data suggest that upper versus lower floral meristem fate in maize is not determined by master regulatory genes, but rather by differences in core physiological processes that coordinate sugar availability, energy homeostasis and plant growth. This foundational work provides important insights into the downstream processes that likely regulate floret abortion and provides a rich set of candidate genes to potentially increase floret fertility in cereal crops and enhance yield.

## Materials and Methods

### Laser Capture Microdissection, RNA isolation and amplification

Ear primorida (1-2 cm) were dissected from greenhouse grown (16 h light at 27°C, 8 h dark at 21°C) B73 plants and immediately fixed and embedded for LCM as previously described (Takacs et al., 2012). Embedded samples were sectioned (8 μm) on a Reichert-Jung (Leica) 2030 rotary microtome and mounted on Zeiss Membrane Slide (1.0 polyethylene naphthalate). LCM was performed using the Zeiss PALM MicroBeam System and a minimum 350,000 μm^2^ tissue/biological replicate for three replicates each of UFM and LFM. Total RNA was extracted using Arcturus PicoPure RNA Isolation kit (Applied Biosystems) and DNase treated using the Qiagen RNase-free DNase set. RNA was amplified (Epicenter TargetAmp 2-Round aRNA Amplification Kit 2.0, Epicentre Biotechnologies), DNase-treated (RapidOut DNA Removal kit, ThermoFisher Scientific) and purified (RNeasy MinElute Cleanup Kit, Qiagen). Quality and size of aRNA was assessed using Bioanalyzer 2100 (Agilent Technologies).

### RNA-seq and data analysis

Library construction (TruSeq Stranded mRNA Sample Prep LS kit) and RNA-seq was performed on an Illumina HiSeq 2500 system by Genomic Sciences Laboratory at North Carolina State University. Raw data was trimmed and low-quality reads were filtered out using trim_galore. Reads were mapped to the maize genome (V3) using Tophat2 (v2.1.0) (Kim et al., 2013) with parameters: --library-type fr-secondstrand --b2-very-sensitive -i 20 and quantified using the htseq-count package with default parameters except: --stranded=yes (Anders et al., 2015). Count tables were analyzed using DESeq2 in the R environment for differential expression analysis (Love et al., 2014). Genes with a minimum read count of 10 in at least two biological replicates, fold change ≥ 2 and adjusted p-value <0.5 were considered differentially expressed. Principal component analysis was performed using DESeq2 (Love et al., 2014) in the R environment and correlation analysis was performed using R package “psych” pairs.panels function.

Gene ontology was analyzed using g:Profiler (Raudvere et al., 2019) with default options except statistical settings: Benjamini-Hochberg FDR and All known genes. Gene enrichment maps were generated using Cytoscape (version 3.7.1) plug-in, Enrichment Map (Merico et al., 2010), with default options except FDR q-value cutoff = 0.05, connectivity = second degree sparse, and size of functional category = 1 to 5000. Functional groups were predicted using MapMan 3.6.0RC1 (X4 annotation), with the maize v3 mapping file (retrieved from Mercator4 Fasta validator with the protein option) (Schwacke et al., 2019). CornCyc 9.0 was used to predict metabolic pathways (Schläpfer et al., 2017). Gene ID description analysis was analyzed with g:Profiler g:Convert functional tab (Raudvere et al., 2019).

### RNA *in situ* hybridization and histochemistry

Inflorescence primordia (1-2 cm) for RNA *in situ* hybridization, lignin/cellulose staining, and immunohistochemistry, were fixed and embedded as described in (Thompson et al., 2009) and sectioned (10 µm) using a Microm HM315 Microtome. Inflorescence primordia for ruthenium red and starch staining were directly frozen with optimal cutting temperature (OCT) embedding medium on dry ice and sectioned (60-80 µm) with a Microm HM550 Cryostat Microtome at -20 °C.

RNA *in situ* hybridization was performed as described previously (Jackson, 1992), with the following modifications. Pronase digestion was performed for 25 minutes at 37 °C; incubated in blocking solution (Sigma Roche) for 1 hour at room temperature before incubation with anti-DIG antibody (1:4000-5000 in blocking solution). After antibody incubation, slides were washed with Buffer A without Triton-X100. Slides were imaged using an Olympus BX-41 compound light microscope and processed using Adobe Photoshop. Probes were generated as described in (Bortiri et al., 2006), using primer sequences listed in Supplemental Table 1.

For pectin, lignin, and cellulose staining, cryosections and rehydrated sections were stained as previously described (Gunawardena et al., 2007; Pradhan Mitra and Loqué, 2014) except staining was performed at room temperature in the dark. Phloroglucinol-HCl-stained samples were immediately imaged with Olympus BX-41 compound light microscope. Calcofluor white/fluorescent brightener 28-stained samples were visualized with an Olympus IX2-DSU Confocal Compound Light Microscope using eDAPI or emDAPI filters. Lugol’s Iodine Solution (Electron Microscopy Sciences) was used to stain starch, washed with 90% isopropanol or ethanol and mounted with histoclear. Maize stem tissue was processed in parallel for all stains.

Immunofluorescence labeling was modified from (Xue et al., 2013). Briefly, rehydrated sections were blocked using 1×PBS with 5% BSA for 30 minutes at room temperature and stained with primary antibodies, LM19 and LM20 (Kerafast, diluted 1:10) overnight at 4°C. After washing in 1×PBS (three washes, five minutes each) sections were incubated for two hours at room temperature with goat anti-Rat IgG (H+L) Alexa Fluor 488 secondary antibody (ThermoFisher Scientific, diluted 1:200). Antibodies were diluted in 1×PBS with 5% BSA and incubated in the dark. Sections were washed three times (five minutes each) with 1×PBS, incubated with 0.02% Toluidine Blue O (1×PBS) for five minutes to quench autofluorescence, and rinsed twice with 1×PBS prior to mounting with antifade medium (Hinnant et al., 2017). Slides were stored at 4°C in the dark prior to imaging with a Zeiss LSM700 laser scanning microscope. Negative controls lacking primary antibody were processed in parallel. Images were processed in Adobe Photoshop and false colored using the Blue Orange icb look up table in ImageJ (Schneider et al., 2012).

## Accession numbers

RNA sequencing data were deposited into NCBI Sequence Read Archive under accession number PRJNA717335.

## Supplemental data

**Supplemental Figure 1.** Cluster and reproducibility analysis of LFM and UFM biological replicates.

**Supplemental Figure 2.** Summary of DEG in the MapMan RNA biosynthesis functional group.

**Supplemental Figure 3.** Summary of DEG in the MapMan phytohormone functional group.

**Supplemental Figure 4.** Summary of DEG in the Mapman protein modification (A) and degradation (B) functional groups.

**Supplemental Figure 5.** RNA *in situ* hybridization of AC217050.4_FG006, *chr101*/GRMZM2G177165, and GRMZM2G101682 in developing inflorescences.

**Supplemental Figure 6.** RNA *in situ* hybridization of *histone H1-like*/GRMZM2G069911 and *XTH9* homolog/GRMZM2G180870 in developing inflorescences.

**Supplemental Figure 7.** RNA *in situ* hybridization of *BBTI*, *adc1* and *pectate lyase* in developing inflorescences.

**Supplemental Figure 8.** Histochemical staining in maize stems and spikelets.

**Supplemental Table 1.** Primers used in this study.

**Supplemental File 1.** Summary of mapping and differential analysis.

**Supplemental File 2.** Summary of Gene Ontology, Mapman, and CornCyc analysis.

## Acknowledgements

We are thankful to Julie Marik, Cathy Herring and Jim Holland for greenhouse and field support, Cindy Kukoly for technical assistance with LCM, Ross Sozzani and Angel de Balaguer for help with RNA seq and early analysis, Katherine Novitzky for technical discussion with RNA *in situ* hybridization, and Allison Beachum, Tom Fink and Elizabeth Ables for help with microscopy. Patrick Horn, Madelaine Bartlett, and Clint Whipple provided useful comments on the manuscript.

## Notes

**Funding information:** This work was supported by grants from the North Carolina Biotechnology Center (#2011-BRG-1213) and the NSF (IOS-1148971).

